# Fast and reliable sgRNA efficiency testing using HIReff

**DOI:** 10.1101/2021.09.15.460502

**Authors:** David Marks, Jannis Anstatt, Lisa Bachmann, Lucia Gallego Villarejo, Alexander Geidies, Thorsten Müller

## Abstract

CRISPR/Cas9 is the method of choice for gene editing like the endogenous knock-in of sequences in order to investigate protein function, abundance or intracellular localization. One of the crucial steps in the preparation of CRISPR/Cas9-mediated knock-ins is the design of sgRNAs, which need to be tested carefully in order to minimize off-target binding and reach highest cleavage efficiency. Usually, sgRNA is evaluated via mismatch cleavage assays, like Surveyor or T7 endonuclease 1 assay. We demonstrate that these methods are often highly cost- and time-intensive with a low sensitivity and high fail rate. As an alternative, we present a new <u>HI</u>TI-based sg<u>R</u>NA <u>eff</u>iciency (HIReff) test to precisely evaluate sgRNA efficiency. HIReff is based on a sophisticated integration vector with on-site generation of a linear donor fragment that allows a comparably easy read-out via fluorescence signal and integrates several internal controls. Next to a quantifiable sgRNA assessment, HIReff provides additional information on the ‘gene/protein to be studied’ abundance, subcellular localization and promoter activity and allows derivation of fluorescence protein labeled clonal lines. We highlight benefits of HIReff in comparison to commonly used enzyme-based assays and demonstrate improved practicability and high sensitivity, while being less time-, labor- and cost-intensive at the same time. Our results suggest HIReff as a fast and easy-to-use alternative for sgRNA efficiency testing.

## Introduction

The method of gene-editing has been revolutionized in the last decade by the development of the CRISPR/Cas9 system. Discovered as a bacterial immune defense mechanism **(Garneau et al., 2010; Horvath & Barrangou, 2010)**, the system has cleverly been adapted as a tool for targeted genome editing and has since evolved to offer an unparalleled set of applications that made gene editing not only more efficient and reliable, but also considerably easier and less time-consuming. One of the major gene editing applications facilitated by CRISPR/Cas9 is the knock-in of genes. The endogenous knock-in of genes offers a broad variety of applications, for instance, it allows to track specific proteins and structures via fluorescent reporter genes, to fuse genes in order to create fusion proteins, or to generate disease models by inserting disease-related gene variants and mutations. These knock-ins take advantage of the doublestrand break (DSB) created through the CRISPR/Cas9 mechanism that triggers the cell’s repair to incorporate a DNA fragment of choice. Different strategies have been devised to achieve a successful knock-in, some of them using the non-homologous end-joining pathway (NHEJ) **(Artegiani et al., 2020; He et al., 2016; Suzuki et al., 2016; Yan et al., 2020)**, while most approaches rely on homology-directed repair (HDR) **(Koch et al., 2018; Roberts et al., 2017; Wang et al., 2015; Yang et al., 2020)**. However, independent of the selected strategy, the key element in every CRISPR/Cas9-mediated knock-in experiment is the single guiding RNA (sgRNA), which determines the target locus and guides the Cas9 nuclease accordingly. Its efficiency and specificity are decisive for the overall efficiency of the knock-in process and it is crucial to carefully design the sgRNAs for every knock-in approach and to have reliable tools at hand to test their efficiency in practice. For the design of sgRNAs, a range of *in silico* tools is available, many of them web-based and freely accessible. These tools greatly facilitate the design of sgRNAs to target specific genes by rating the quality of possible sgRNA candidates based on different criteria and input data in order to minimize off-target effects **(Cui, Xu, Cheng, Liao, & Peng, 2018)**. Examples for popular web tools are CHOPCHOP (https://chopchop.cbu.uib.no/, **(Montague, Cruz, Gagnon, Church, & Valen, 2014)**), CasFinder (https://arep.med.harvard.edu/CasFinder/, **(Aach, Mali, & Church, 2014)**), or E-CRISP (http://www.e-crisp.org/E-CRISP/designcrispr.html, **(Heigwer, Kerr, & Boutros, 2014)**). After successful identification and design, the suitable sgRNA candidates require practical testing for specificity and cleavage efficiency. The method of choice is usually a mismatch cleavage assay, which relies on the error susceptibility of the cell’s repair machinery **(Qiu et al., 2004; Vouillot, Thélie, & Pollet, 2015)**. Briefly, transfection with sgRNA and Cas9 leads to the induction of short heterogenous insertions and deletions (InDels) at the site of the DSB resulting in heterogeneous cell populations comprising a mix of mutant and wild-type cells which is the basis of mismatch cleavage assays using commercially available endonucleases, such as Surveyor^™^ **(Qiu et al., 2004)** or T7 endonuclease 1 (T7E1). Generally, to perform these assays, the DNA fragments are denatured into single strands, and then re-hybridized allowing annealing of mutant and wild type strands. Mismatch-specific endonucleases induce cleavage of heteroduplexes, so that subsequent agarose gel electrophoresis analysis reveals two shorter cleavage products besides the full-length fragment, indicating that the sgRNA successfully initiated a DSB and the ensuing NHEJ repair pathway causing indels in the target region. The ratio of wild type to cleavage product serves as an indicator for the relative sgRNA efficiency. While these kinds of assays are advertised as fast and simple methods, we have often had contrary experiences, where besides being highly cost- and time-intensive, these assays were limited in detection sensitivity and efficiency, especially in stem cell systems such as induced pluripotent stem cells (iPSCs) due to low transfection efficiencies **(Ardehali et al., 2011; Ihry et al., 2018)**.

Here, we report an alternative evaluation system for sgRNA efficiency including fluorescence readout and information about promoter activity: the homologous-independent targeted integration (<u>HI</u>TI)-based sg<u>R</u>NA <u>eff</u>iciency test (HIReff). HIReff enables an endogenous promoter-dependent mNectarine reporter gene knock-in allowing direct and easy confirmation of sgRNA functionality by detection of fluorescence signals.

## Materials and Methods

### Cell culture and transfection

HEK293T were routinely cultured in Dulbecco’s Modified Eagle Medium (DMEM, Sigma-Aldrich) with 10 % FBS, 1 % Pen/Strep and 1 % L-glutamine on 100 mm culture dishes, T-75 culture flasks or 6-, 12-well plates. Transfection was performed using the K2^®^ Transfection system (Biontex, Germany) according to the manufacturer’s instructions with minor modifications. To test the transfection and CRISPR/Cas9-mediated insertion efficiency, HEK293T cells were transfected with HIReff Integration Plasmid, HIReff sgRNA A, HIReff sgRNA B and Cas9 expressing vector. Results were monitored by fluorescence microscopy and DNA analysis and sequencing.

### Surveyor assay

To test the activity of four different Cas9 constructs and two sgRNAs targeting the Fe65 gene, a surveyor assay was performed.The sgRNAs were designed and cloned according to Kwart et al. **(Kwart, Paquet, Teo, & Tessier-Lavigne, 2017)**. HEK293T cells were transfected with a combination of one Cas9 construct (pCas9_GFP, pCas9_DsRed, pCas9_Neomycin or pCas9_Blasticidin) and either *sgRNA_Fe65_1 or sgRNA_Fe65_2* (targeting the C-terminal domain of Fe65). To detect specific cleavage points by Cas9, genomic DNA from transfected HEK293T cells was isolated using the NucleoSpin^®^ DNA RapidLyse Kit (Machery-Nagel, Germany) and the genomic regions were amplified by PCR using primers (Surveyor_Fe65_for CTGCGTGCATGGTAAGCTACTAGTAGG and Surveyor_Fe65_rev CACCTGAAACACATGGGGATGGAGA). Presence of Indels in the PCR product were tested using the Surveyor^®^ Mutation Detection Kit (Integrated DNA Technologies) according to the manufacturer’s instructions and the final product was analyzed on a 2 % agarose gel. DNA from non-transfected HEK293T cells was identically obtained and analyzed as a control. Additionally, two standard fragments with (GC) and without (C) mutations were analyzed as controls following Surveyor^®^ Mutation Detection Kit (Integrated DNA Technologies) manufacturer’s protocol.

### Establishment of HIReff assay

Plasmid digestion using BspEI, HF-BamHI, HF-EcoRI, HF-AgeI and BsmBI (New England Biolabs) as well as dephosphorylation using antarctic phosphatase (New England Biolabs) was performed according to the manufacturer’s instructions. PCRs were performed using the Q5^®^ High-Fidelity DNA Polymerase kit (New England Biolabs) according to manufacturer’s instructions. PCR products were analyzed via agarose gel electrophoresis on a 1 % (w/v) agarose gel with 0.001 % GelGreen^®^ Nucleic Acid Gel Dye (Biotium) (90 V, 30 min). Samples were priorly loaded using Gel Loading Dye, Purple (6X, New England Biolabs). After separation, DNA fragments were extracted from the gel and purified using the NucleoSpin^®^ Gel and PCR Clean-up kit (Machery-Nagel). Sticky ends were ligated using the T4 ligase (New England Biolabs) according to the manufacturer’s instructions. In-Fusion cloning was performed using the In-Fusion^®^ HD Cloning Plus Kit (Takara Bio, Japan) following the manufacturer’s protocol. Subsequently, the constructs were transformed into competent DH5α E. coli cells according to Addgene protocol (https://www.addgene.org/protocols/bacterial-transformation/?hypothesisAnnotationId=XWgbiMsUEees2s_t2EHX6A), followed by plasmid DNA isolation using the Monarch^®^ Plasmid Minprep kit (New England Biolabs) or a NucleoBond^®^ Xtra Midi kit (Machery-Nagel). DNA identity was confirmed using Sanger sequencing.

### Testing the HIReff assay

The HIReff plasmid mix, consisting of the HIReff Integration Plasmid, HIReff sgRNA A, HIReff sgRNA B, Cas9 and sgRNA GOI was used for transfection of HEK293T cells using the K2^®^ Transfection System (Biontex). To test the efficiency of the sgRNAs and to verify the insertion of the plasmids in the host genome, HEK293T cells were analyzed by fluorescence microscopy 24 and 48 h after transfection regarding the mNectarine and EGFP signal. HEK293T cells were analyzed via confocal microscopy and afterwards cells were harvested, and the genomic DNA was isolated using the NucleoSpin^®^ DNA RapidLyse Kit (Machery-Nagel). Genomic DNA was amplified using primers HIReff Insert verification For (CTGCGTGCATGGTAAGCTACTAGTAGG, identical to Surveyor_Fe65_for) and HIReff Insert verification Rev (CCGGTGAACAGCTCCTCGC). While HIReff Insert verification For is a primer designed to bind in the host genome to verify the correct insert, the HIReff Insert verification Rev is able to bind reverse in the construct inserted. Fragments were then analyzed on a 1 % agarose gel and the band with the expected fragment size was isolated with the NucleoSpin^®^ Gel and PCR Clean-up Kit (Machery-Nagel, Germany). Results were validated by Sanger sequencing in order to confirm the correct insertion. Alternatively, after 14 days when BFP signal vanished, EGFP positive cell colonies were picked and further cultured to generate a stable clonal cell line with a mNectarine cassette fused at the C-terminal domain of the Fe65 protein.

### Microscopy

To determine the transfection efficiency, transfected HEK293T were imaged with a Olympus IX51 inverted microscope system equipped with a 10, 20, and 40x magnification objective. Pictures were obtained using excitation wavelengths of 360-370 nm (BFP), 450-480 nm (GFP) and 520-550 nm (mNectarine) and emission wavelengths of 400-420 nm (BFP), 500-520 nm (GFP) and 565-575 nm (mNectarine). For confocal imaging, the Leica TCS SP8 confocal microscope system was used. Images were recorded with a 100x oil objective (1.4 NA) using excitation wavelengths of 405 nm (BFP), 488 nm (GFP) and 587 nm (mNectarine) and emission wavelengths of 410-483 (BFP), 493-592 nm (GFP) and 592-779 nm (mNectarine). Images were acquired in a sequential scan at scan speed of 400 Hz and a line average of 16 into ca. 17 pixels/μm resolution images. Hybrid detectors (HyD) were used for BFP/mNectarine and a photomultiplier tube detector (PMT) for GFP.

## Results

### Surveyor assay for testing sgRNA efficiency demonstrates limited sensitivity

One of the most widespread methods to test the activity of Cas9 constructs or the efficiency of sgRNAs is the commercially available Surveyor assay. It aims at detecting genetic heterogeneity in cell culture caused by the error-prone repair of Cas9-induced ds-breaks by non-homologous end joining (NHEJ) (Figure 1A). The presence of genetic heterogeneity, shown by cleavage of DNA fragments with a mismatch at the cutting site by the surveyor nuclease, indicates a functional Cas9 enzyme and active sgRNA constructs, whereas the absence of cleavage products indicates problems at some point in the genome editing process. The whole assay includes several steps including DNA isolation, PCR, denaturation/annealing and gel imaging (Figure 1B). To analyze the performance of the Surveyor assay, we investigated the activity of two different sgRNAs in combination with four different Cas9 constructs (Figure 1C). As a target we chose Fe65, a protein with elementary function in Alzheimer’s disease, which is in the focus of our research projects **(Kolbe et al., 2016; Marks et al., 2021; Nensa et al., 2014)**. For all combinations as well as the negative control (no sgRNAs), the band corresponding to the uncut PCR product (yellow arrow) was by far the most intense whereas signals corresponding to the cut PCR products (red arrows) were difficult to discern or sometimes absent, even though these sgRNAs and Cas9 enzymes were successfully employed for genome editing in other experiments (not shown). As control, DNA from non-transfected (wt) cells as well as synthetic DNA oligonucleotides with GC mismatch or with C (no mismatch) were used. It is apparent that even with these synthetic controls the band corresponding to the cleavage product is only clearly visible when the gel is excessively loaded with DNA. As a consequence, we evaluated the assay as less solid and aimed to establish a more reliable approach.

**Figure 1.**
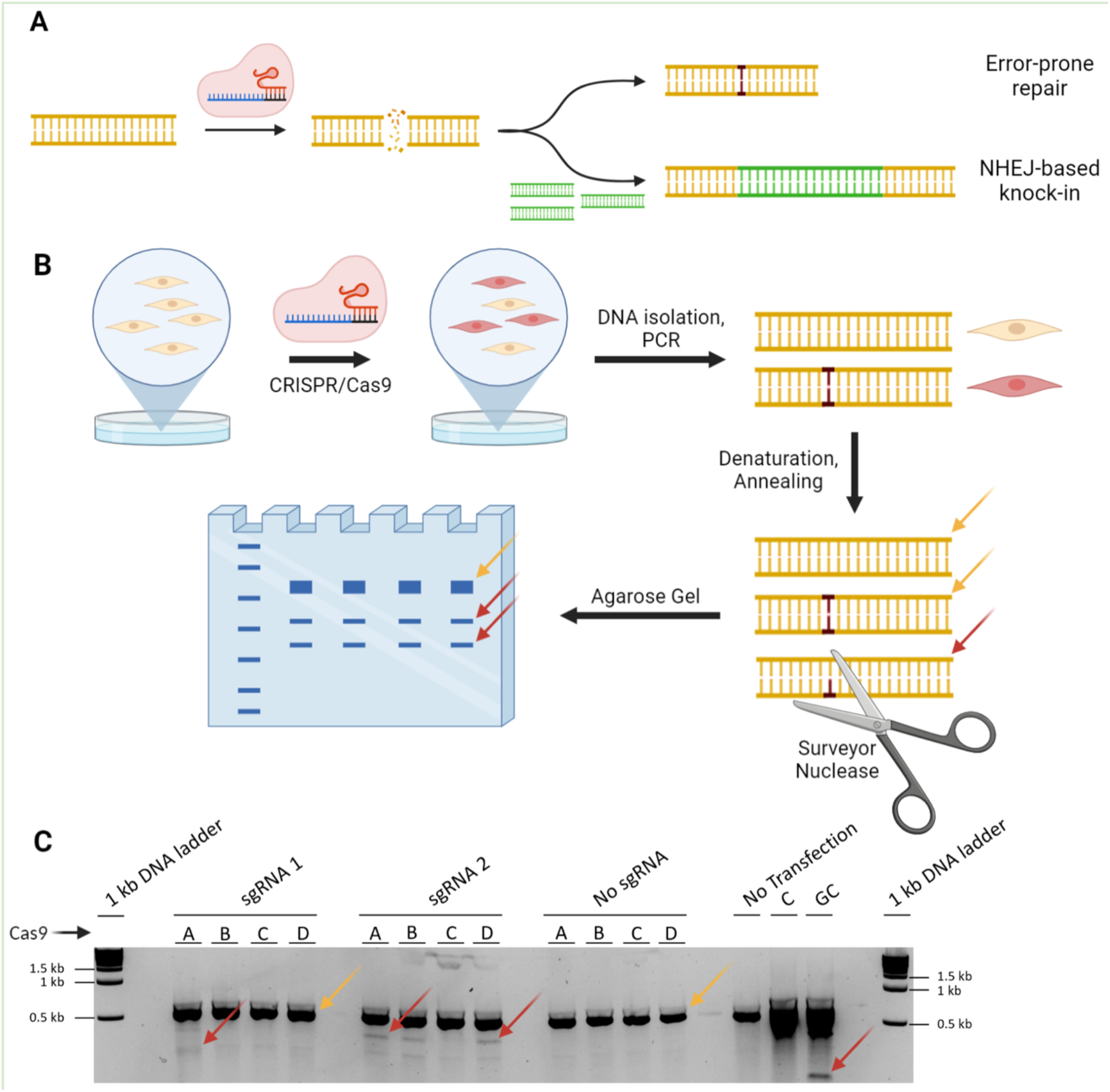
Overview of non-homologous end joining and surveyor assay. **(A)** Schematic representation of non-homologous end joining (NHEJ): The genomic DNA is cut at a specific place by a Cas9/sgRNA-complex. The resulting double strand break (DSB) can be repaired by error-prone NHEJ. In the presence of an excess of additional DNA fragments, those can be incorporated into the genomic DNA, resulting in NHEJ-based knock-in. **(B)** Schematic representation of the surveyor assay: Cells are transfected with Cas9 and a sgRNA, resulting in DSB which are repaired by NHEJ, frequently disrupting the target gene. Genomic DNA isolation and PCR amplification of the target gene yields DNA fragments with or without mutation (Insertions or Deletions). Denaturation and annealing of the DNA fragments produces duplexes with (red arrow) or without (yellow arrow) mismatches. Only fragments with mismatches are cut by the surveyor nuclease. Fragment lengths are detected on an agarose gel. **(C)** Example result of a surveyor assay. Three different sgRNAs were tested in combination with four different Cas9 constructs. As control, genomic DNA from nontransfected cells and DNA fragments with (C) or without (GC) mismatch were utilized.

### Establishment of the <u>HI</u>TI-based sg<u>R</u>NA <u>eff</u>iciency (HIReff) assay

Our central goal was the establishment of a reliable and easy to use approach combined with a fast read-out for sgRNAs addressing mRNA sequences. The new HIReff assay enables this readout by fluorescence microscopy and opens up the possibility for the generation of clonal lines with C-terminally tagged fusion proteins. For this, the HIReff integration plasmid is transfected together with constructs containing Cas9, two HIReff sgRNAs A/B and the to be evaluated sgRNA for the gene of interest (GOI, Figure 2A). As a consequence of the design, two DSB will be set on the integration vector within the cellular nucleus providing an on-site linear knock-in fragment. The HIReff integration plasmid consists of multiple fluorescent transfection and integration controls to rapidly visualize results. Upon transfection, the BFP cassette serves as transfection control and shows a transient blue fluorescence. The cellular NHEJ repair mechanism, caused by a successful induced double strand break of the sgRNA for the GOI will integrate the linearized HIReff integration cassette (Figure 2B). This cassette consists of a mNectarine sequence, which requires an endogenous promoter and enables the creation of C-terminal tagged fusion proteins, if inserted correctly. As a consequence, next to successful sgRNA mediated cleavage, promoter activity can be directly assessed using the mNectarine fluorescent protein. Additionally, an EGFP/BleoR cassette is inserted, which is the second readout of the tested sgRNA since this cassette is driven by its own strongly expressing promotor. Since the readout is fluorescent based, the cells can be observed after a short period of time, thus enabling a fast readout system to evaluate the used sgRNA. Optionally, the sgRNA can be designed to enable the generation of a fusion-protein at the C-terminal domain of the GOI. The red positive cells can then be sorted via FACS to create a stable cell line (Figure 2C). The correct integration can be checked via PCR and Sanger sequencing.

**Figure 2.**
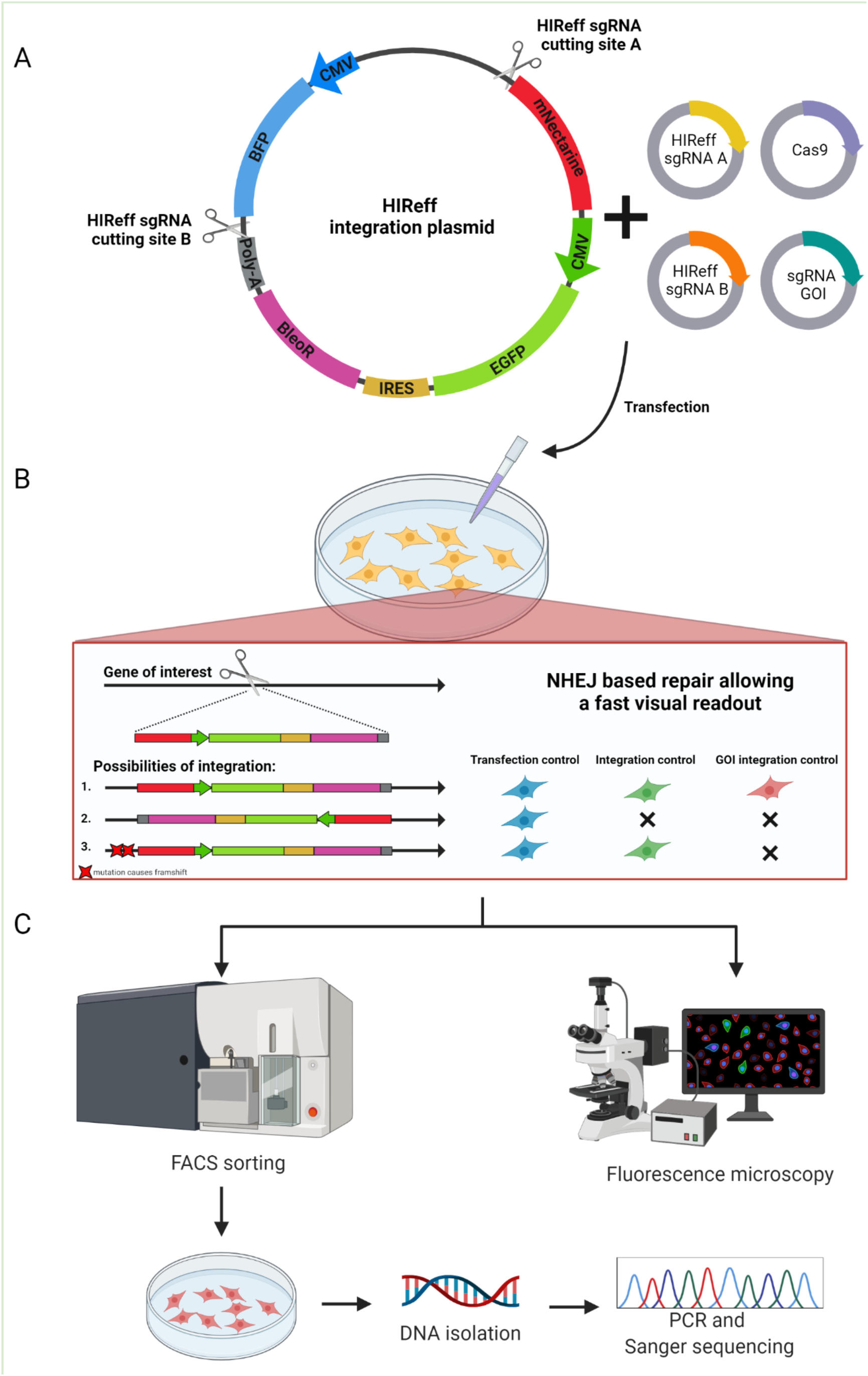
HIReff assay design and readout. **(A)** Schematic overview of the HIReff integration plasmid and the transfection mix consisting of two HIReff sgRNAs cutting site A and B, Cas9 and the sgRNA GOI, which is to be evaluated. The HIReff integration plasmid consists of three fluorescent elements: A BFP cassette, serving as a transfection control, mNectarine without promoter for the creation of C-terminal fusion proteins (and thereby validating sgRNA dependent DSB) with the GOI and an EGFP/BleoR cassette with a strong independent CMV promoter to visualize successful and stable integration. **(B)** Upon transfection, the HIReff integration plasmid will be linearized and integrated into the gene of interest via the NHEJ repair, caused by the sgRNA mediated DSB for the GOI. This can lead to different possible integration of the linearized cassette: (1) the cassette is correctly inserted, leading to the expression of all three fluorophores. (2) the cassette is reversed inserted, thus inhibiting the expression of the green and red fluorophore. Only the transfection control in blue is visible. (3) the cassette is integrated but the error-prone NHEJ repair mechanism leads to mutation-caused frameshifts, leading to an unsuccessful expression of the red fluorophore. Since the EGFP and BleoR are driven by their own promoter, the green fluorescence is still visible. **(C)** The transfected cells can either be evaluated with fluorescence microscopy, evaluating the success of the used sgRNA of the GOI or the cells can be further used when a fusion protein with the GOI is desired. Thus, the cells can be FACS sorted for the red fluorescence or treated with bleomycin. Upon DNA isolation, the successful integration of the desired fusion protein is evaluated with PCR and Sanger sequencing.

### Application of HIReff to FE65 revealed fast sgRNA testing and generation of fusion proteins

As a next step, we performed the practical test of our HIReff approach for sgRNA Fe65 as gene of interest (GOI) that is supposed to induce a CRISPR/Cas9 DSB at the C-terminal end of the Fe65 gene. Accordingly, we aimed to test both features of HIReff, the sgRNA test associated with the generation of an endogenous fluorescent protein tag. According to other pre-work **(Schrötter et al., 2013)**, successful HIReff should result in a punctuated cytosolic localization of Fe65. We transfected HEK293T cells with HIReff Integration Plasmid, HIReff sgRNA A, HIReff sgRNA B, sgRNA Fe65 (GOI) and pCas9-Blasticidin, and analyzed the fluorescence signals via microscopy imaging 48 h after transfection in order to determine transfection and, finally, cleavage efficiency (Figure 3A). Expectedly, we observed a greater quantity of GFP-positive cells than mNectarine-positive cells (GFP 51.7 ± 4.3 %; mNectarine 15.7 ± 2.3 %; Figure 3B). As GFP is independent of co-expression under the Fe65 promoter, it can be considered an indicator for transfection efficiency directly after transfection, since its CMV promoter ensures expression with or without integration. Compared to mNectarine expression, which is dependent on the endogenous Fe65 promoter, and therefore an indicator for correct insertion, we can estimate that in approximately 30.4 % of transfected cells, the insertion cassette was incorporated successfully. Additionally, transfection was performed omitting one of the sgRNAs, respectively, which revealed strong BFP and GFP signals, thus effective transfection, but no specific signals for mNectarine resulting from integration (Figure 3C) which strongly emphasizes the specificity of the approach. For closer investigation of subcellular localization, we performed confocal live cell imaging, which showed that transfected cells exhibited mostly uniform BFP and GFP signals (Figure 3D). Interestingly, mNectarine signals were organized in spheres distributed throughout the cytoplasm corresponding to the more specific Fe65 localization (Figure 3E). Sanger sequencing of the knock-in region confirmed correct integration of the fragment (not shown). Altogether, the HIReff approach resulted in the expected fluorescence signal read-out which gave clear evidence of sgRNA specificity and efficiency as compared to transfection efficiency.

**Figure 3.**
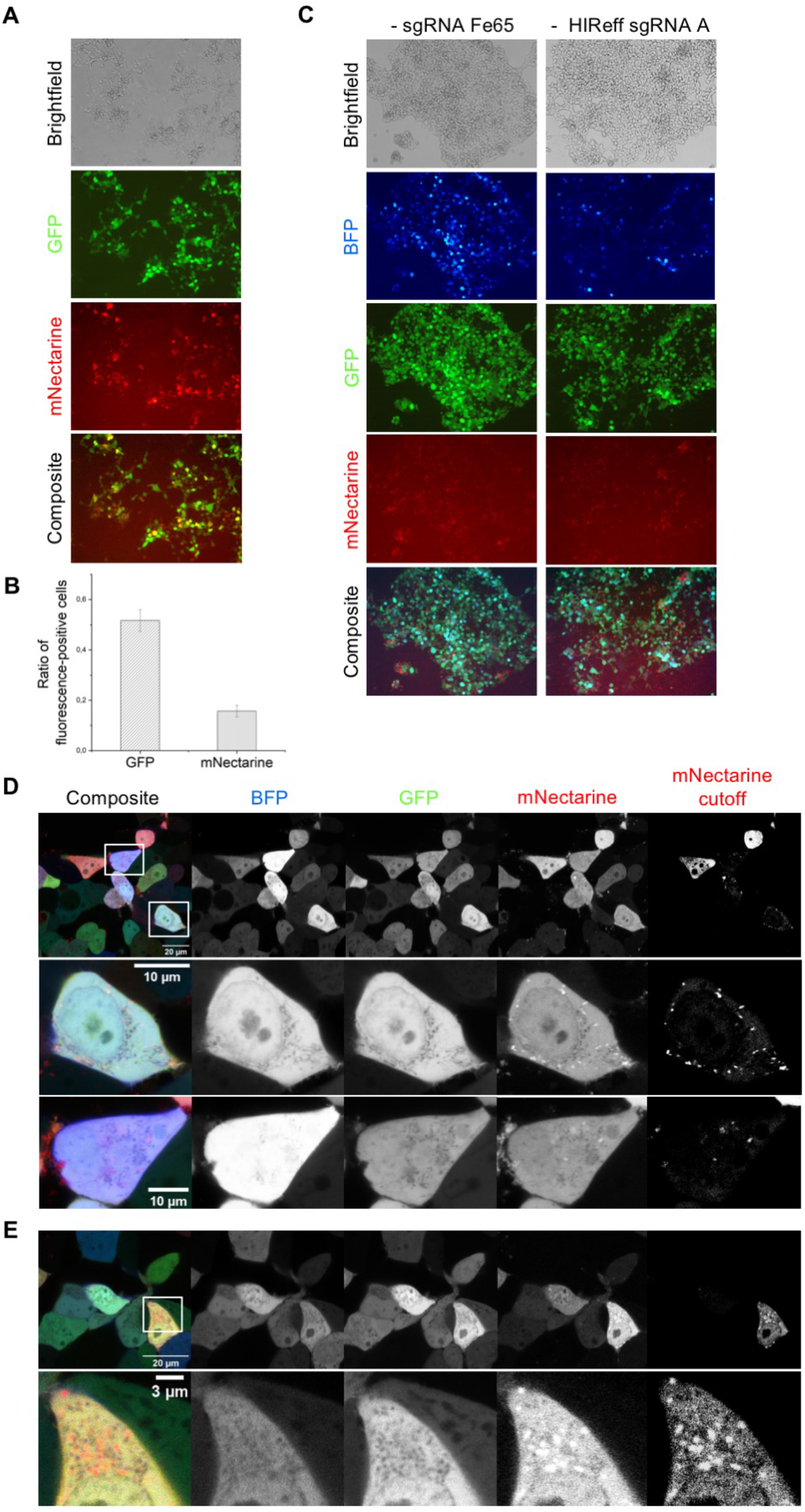
HIReff in practice - Fluorescence readout of sgRNA cleavage efficiency. **(A)** After transfection with HIReff Integration Plasmid, HIReff sgRNA A, HIReff sgRNA B, sgRNA Fe65 and pCas9-Blasticidin, HEK293T cells were analyzed via fluorescence microscopy (20x magnification). Shown are exemplary images of GFP (green) and mNectarine (red) signals next to a brightfield image and an overlay (composite) demonstrating successful transfection (GFP) and incorporation of the knock-in fragment (mNectarine). **(B)** GFP and mNectarine signals were analyzed quantitatively as ratio of fluorescent cells relative to the total amount of cells, depicted in a bar chart. Error bars represent standard errors. **(C)** Cells were transfected and analyzed analogous to (A), but sgRNA Fe65 or HIReff sgRNA A were omitted, respectively. Shown are BFP (blue), GFP (green), mNectarine (red) signals as imaged via fluorescence microscopy. **(D),(E)** Transfected cells were additionally analyzed via confocal microscopy offering higher resolution of cellular localizations. Shown are exemplary images of BFP (blue), GFP (green), mNectarine (red) signals, and an overlay of all channels (composite). In addition, signal intensity was edited subsequently to enhance mNectarine signals (mNectarine cutoff). Specific cells are each shown in a close-up view as indicated by boxes. Cells exhibit uniform BFP signals, while mNectarine signals are spheres distributed throughout the cells.

## Discussion

Efficient sgRNA-directed Cas9 cleavage is a central requirement for reliable gene editing. The commonly used enzyme-based methods to analyze sgRNA-dependent cleavage potential like the Surveyor-assay have major disadvantages due to low robustness, poor signal intensities, relatively high DNA amounts needed for analysis, and time required. Moreover, in our hands, several replicates were often needed to get sufficient results showing (highly fluctuant) band intensities to enable a verified read-out of sgRNA functionality. Enzyme-based assays particularly rely on a high transfection efficiency. The comparatively low sensitivity of agarose gels makes it difficult to detect the cleavage fragments if only a few cells were successfully edited due to low transfection efficiency. As the production of DNA fragments of the right size is the only read-out of the surveyor assay it is impossible to quickly determine whether one of the CRISPR/Cas9 components was non-functional or whether the assay failed due to low transfection efficiency or an inactive surveyor nuclease. These limitations exacerbate quantification of the DNA cleavage by the Cas9/sgRNA complex and thereby assessing quality of the selected sgRNA. Consequently, there is a need for an alternative tool, which is easy to use and delivers reliable results. Within this work, we present HIReff, a new method based on the NHEJ-driven integration of a donor fragment at the DNA DSB induced by an sgRNA/Cas9 complex to be tested. Due to the HITI-driven highly efficient generation of the linear donor fragment within the cell, HIReff is highly efficient for fast read-out, which is accomplished by the (promoter-free) integration of a mNectarine fluorescent protein sequence into the target genome. As a consequence, sgRNA/Cas9 cleavage efficiency is evaluable at ~24 h after application of HIReff by fluorescence microscopy and quantifiable by the ratio of mNectarine+ to all cells. The fast read-out enables efficient CRISPR gene editing optimization, e.g. the comparison of newly engineered Cas9 enzymes or sgRNA concentrations. In contrast, enzyme-based assays include multiple steps including transfection, genomic DNA preparation, PCR, hybridization, endonuclease digestion, and DNA gel electrophoresis being a time-consuming, labor-intensive and less reliable approach as demonstrated in this work. As a consequence of this long workflow, for those combinations where the enzyme-based assay shows negative results it cannot be excluded that the negative result was caused by e.g. issues with the transfection procedure or hybridization, although the used sgRNA had high efficiency. In contrast, a one-step transfection and standard fluorescence microscopy, available in virtually all cell/molecular biology labs is sufficient for sgRNA analysis by HIReff. Next to sgRNA assessment, HIReff highlights additional features for the gene of interest. As a result of the integration vector design, promoter activity of the gene to be studied can be estimated by the mNectarine fluorescent protein intensity and in case of sgRNAs addressing the C-terminal GOI end, the subcellular localization of the protein under endogenous expression conditions is provided as shown for FE65 in this work. HIReff offers internal control of the approach by additional modules included in the integration vector. This includes a CMV-BFP cassette in order to control transfection efficiency. For sgRNAs addressing non or low expressed promoters or non mRNA targets, an additional CMV-GFP cassette was designed to be inserted into the target site. As a consequence, successful integration and thereby sgRNA functionality is evaluable as permanent positive GFP fluorescence while weakening BFP signal over time. While HIReff was primarily designed as a sgRNA test tool, it is usable for the establishment of fluorescence protein labeled cell lines as well. Integration of a Bleomycin resistance gene enables efficient screening for cells of interest, which is further engageable by FACS separation. Researchers aiming to use HIReff in this direction, need to take notice that triplet InDels might occur due to the error-prone NHEJ, which is used as integration strategy, and accordingly, sequencing of the fusion protein sequence is mandatory, which is, anyway, part of best lab practice. Testing for proper integration on the genomic level is further facilitated by HIReff as PCR amplifications of the region of interest will directly show a successful knock-in in consequence of the large knocked-in fragment causing a band shift on the agarose gel. As a side note, usage of NHEJ in HIReff enables applicability in the majority of cell lines, as this DSB repair method is generally favored to HDR **(Devkota, 2018; Ghezraoui et al., 2014; Jasin & Rothstein, 2013; Shrivastav, De Haro, & Nickoloff, 2008)**. Being a highly efficient tool in our hands, we need to point out that further optimization could strengthen HIReff even more, e.g. the joining of the multiple vectors or an AAV-driven system for the editing of cell lines known to be hard to transfect by standard methods. Next to the illustrated lab-associated advantages, HIReff is of interest from the economic point of view taking the high costs for commercial enzyme-based kits into account. Thus, for labs working in gene editing, HIReff offers an interesting alternative for fast, reliable, and cost-effective sgRNA testing providing additional results on promoter activity, protein of interest abundance or subcellular localization and fostering establishment of fluorescence labeled clonal lines.

## Acknowledgements

This work was supported by funding from Deutsche Forschungsgemeinschaft (DFG, MU3525/3-2), Mercur (Pr-2016-0010) and the Federal Ministry of Education and Research Germany (Bundesministerium für Bildung und Forschung; BMBF) (OrganSARS, 01KI2058). The instrument Leica TCS SP8 was supported by an instrument grant from the German Research Foundation (INST 213/886-1 FUGG). Figures in this paper were created with BioRender.com.

